# An Advanced Framework for Time-lapse Microscopy Image Analysis

**DOI:** 10.1101/2020.09.21.303800

**Authors:** Qibing Jiang, Praneeth Sudalagunta, Mark B. Meads, Khandakar Tanvir Ahmed, Tara Rutkowski, Ken Shain, Ariosto S. Silva, Wei Zhang

## Abstract

Time-lapse microscopy is a powerful technique that generates large volumes of image-based information to quantify the behaviors of cell populations. This method has been applied to cancer studies to estimate the drug response for precision medicine and has great potential to address inter-patient (or intertumoral) heterogeneity. A couple of algorithms exist to analyze time-lapse microscopy images; however, most deal with very high-resolution images involving few cells (typically cell lines). There are currently no advanced and efficient computational frameworks available to process large-scale time-lapse microscopy imaging data to estimate patient-specific response to therapy based on a large population of primary cells. In this paper, we propose a robust and user-friendly pipeline to preprocess the images and track the behaviors of thousands of cancer cells simultaneously for a better drug response prediction of cancer patients.

**Availability and Implementation:** Source code is available at: https://github.com/CompbioLabUCF/CellTrack

**ACM Reference Format:** Qibing Jiang, Praneeth Sudalagunta, Mark B. Meads, Khandakar Tanvir Ahmed, Tara Rutkowski, Ken Shain, Ariosto S. Silva, and Wei Zhang. 2020. An Advanced Framework for Time-lapse Microscopy Image Analysis. In *Proceedings of BioKDD: 19th International Workshop on Data Mining In Bioinformatics (BioKDD).* ACM, New York, NY, USA, 8 pages. https://doi.org/10.1145/nnnnnnn.nnnnnnn

## 1 INTRODUCTION

An increasing number of biological and biomedical studies are using time-lapse microscopy imaging data to observe the dynamic behavior of cells over time [2, 16, 17]. Such imaging platform quantifies the number of cells and their sizes, shapes, and dynamic interactions across time [16]. These quantitative properties provide critical insight into the fundamental nature of cellular function.

Cancer cells vary widely in their response to drug treatment, drug tolerance development, survival, and metastatic potential [17, 20, 26]. Recent cancer studies relied on time-lapse microscopy imaging data to estimate the chemosensitivity of patient-derived cancer cells based on their quantitative properties. Such imaging platforms aid in characterizing tumor heterogeneity by creating sub-populations with different degrees of chemosensitivity to a given drug [19–21]. Figure 1 presents the microscopy image for a one-time point. Most of the white spots in this image represent cells. Some of them are tumor cells (alive or dead) and some of them are macrophages. The green rectangle in Figure 1 points out some of the stromal cells, which are the translucent white. In this study, the time-lapse microscopy imaging data imaged thousands of primary tumor cells every 30 minutes for six days with 289 time points (289 images) based on our EMMA platform (ex vivo imaging platform) [19, 20]. The 289 images can be stacked together to generate a time-lapse video that contains the necessary information to estimate the effectiveness of a drug in a cancer patient at a specific dose. As time goes by, some tumor cells undergo cell death due to the action of the drug, while some are attacked by macrophages. An example of this is shown in the yellow rectangle in Figure 1 which presents one macrophage ‘eating’ a tumor cell (i.e., phagocytosis) as they come in contact with each other.

**Figure 1:**
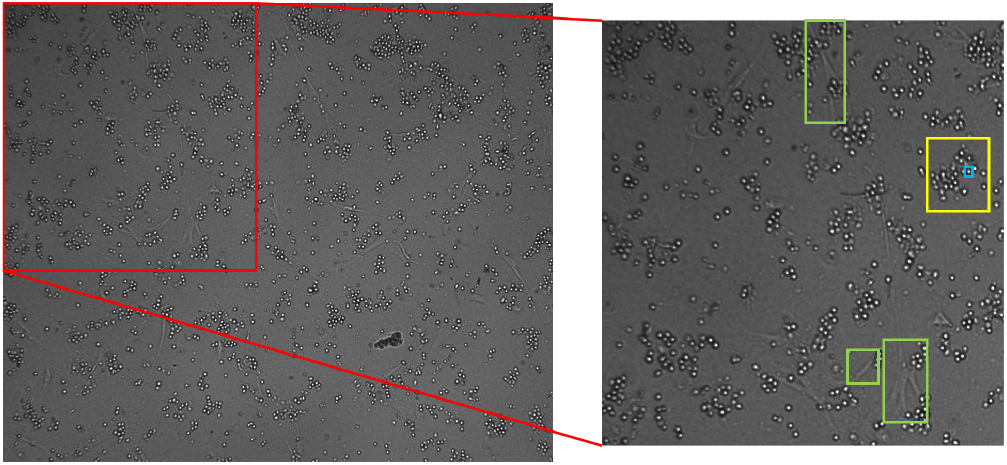
Time-lapse microscopy image. On the left is a microscopy image with an enlarged portion displayed on the right. The white spots represent cells, some of which are macrophages. The blue rectangle shows a macrophage that is devouring the cells in the yellow rectangle. The translucent white lines in the green rectangles are stromal cells.

There are a few algorithms available to analyze time-lapse microscopy imaging data [3–6, 10, 14, 15, 25, 27]. Most work on the images involving very few cells (typically cell lines) at a very high-resolution, making cell segmentation and tracking relatively straight-forward. Unlike most studies, we are working with microscopy imaging data that captures several thousand primary cells (derived from patients) in every frame. Due to the massive number of cells, the resolution at the cellular scale is relatively weak, making cell detection and tracking a problematic task. To overcome the challenges and create a better prediction of the cancer patients’ drug sensitivities, we developed an open-source time-lapse microscopy imaging data analysis pipeline. This pipeline computes differences in sequential microscopy images so that it can track cells, classify live cells, detect cell death, identify cells based on behavior, and track phagocytosis events.

## 2 METHODS

In this section, we introduce each step of our proposed five-step framework for time-lapse microscopy imaging data analysis. The first three steps are image preprocessing, cell detection, and cell tracking. After the tumor cells are successfully detected and tracked, a classification algorithm is applied to distinguish the live and dead cells at each time point (i.e., image). Also, the framework can detect events of phagocytosis (i.e., macrophages devouring tumor cells) by applying a density-based spatial clustering algorithm.

### 2.1 Image Pre-processing

#### 2.1.1 Image Luminance Adjustment

Our framework is invariant to the three channels (i.e., red, green, blue) of the original RGB images, with all three channels having the same values for each pixel. To avoid the redundancy of performing the same operation to each of the three channels separately, we convert the RGB images into gray scale and use these gray scale images for all following analyses.

In this study, we use luminance values to detect cells and track their behaviors in the images; however, as can be seen in Figure 2, the luminance values of the converted images at two-time points (*a* and *b*) can be different. The luminances of the converted images are not in the same scale nor under the same distribution. This difference in luminance adversely affects the cell detection and tracking accuracy of our model; therefore, we need to make the images compatible in terms of luminance.

**Figure 2:**
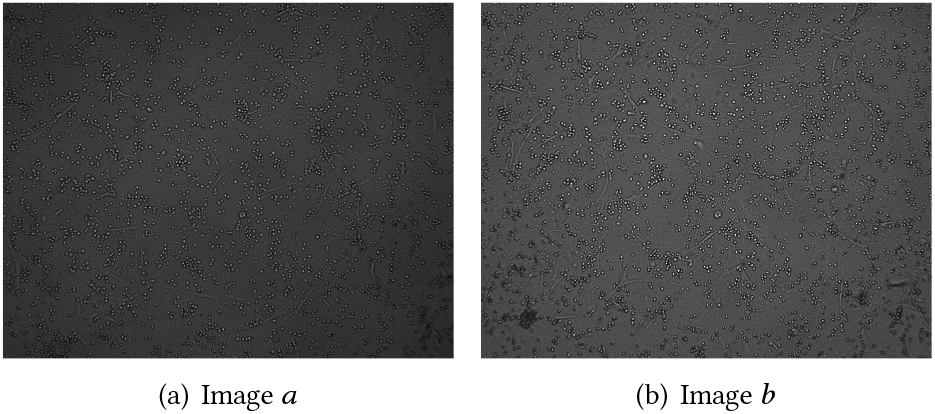
Original Images.

To adjust the luminance of the images, We use Image Histogram Sliding (IHS) [11]. Figure 3 shows the image histograms before and after IHS. In the image histograms, the horizontal axis is the pixel gray level values, while the vertical axis is the frequency of those values. The most frequent pixel values in image *a* and image *b* are 56 and 79, respectively. 56 occurs 73122 times in image *a* and 79 occurs 65332 in image *b*. These create the two peaks in Figure 3 (a). To make the two images have similar brightness, we shift the peak pixel values of both images to a target peak value (100 in this paper). For example, in image *a*, all pixels with value 56 will shift to 100, the rest of the pixel values changing accordingly. Similarly, in image *b*, all pixels at value 79 will be moved to 100 and the rest of the values will be scaled accordingly. The following equation can describe this process:

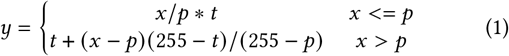

where *x* is an arbitrary pixel value in the original image, and *y* is the generated pixel in the new image. *p* is the peak pixel in each image. *t* is the target peak value.

**Figure 3:**
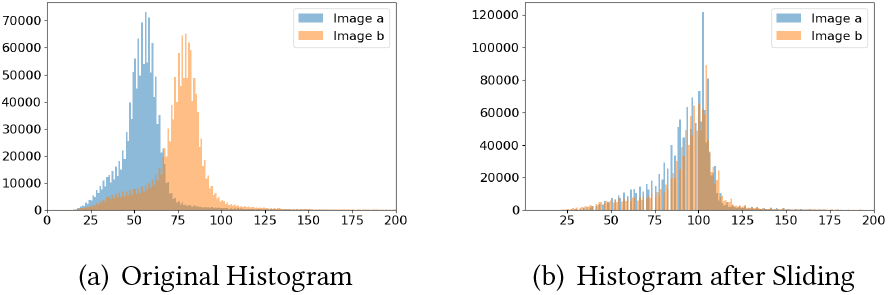
The Histograms of Image *a* and Image *b*.

The IHS adjustment produces new histograms displayed in Figure 3 (b), which shows the histograms almost completely overlapping. Thus, by using IHS, we can regenerate image *a* and image *b* while compensating for the difference in luminosity between frames, as shown in Figure 4. In our analysis, all images align to the same target peak value by applying IHS.

**Figure 4:**
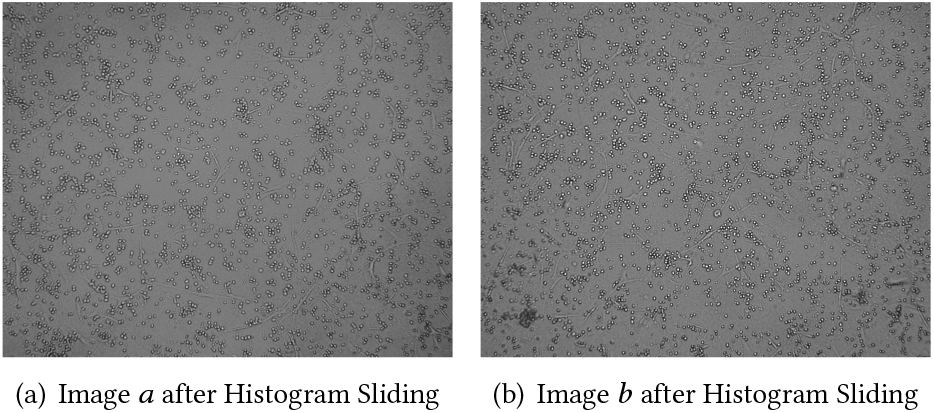
Images after Histogram Sliding.

#### 2.1.2 Image Scaling

The resolution of the original image is 1328 × 1048, whereas, for one cell, it is approximately 10 × 10. This resolution, at the individual cellular scale, is rather low. Therefore, if we set a threshold pixel value to extract the white core or black edge of the cell, there will not be enough pixels representing a cell for further analysis. For example, if we set a pixel value threshold of 130 to filter out the dark pixels from the original image shown in Figure 5 (a), we would end up with Figure 5 (b): very few white pixels representing a cell. The shape of the cell in that image is defined arbitrarily by limited pixels and does not illustrate the actual structure. If one more pixel were to be filtered out using a higher threshold, the shape would change drastically. It is of paramount importance that we have a well-defined form for each cell since our model works based on the detection of the shape of the cells. As a result, to get a smooth contour of the cell, we use image interpolation [23] to improve the resolution of the images.

**Figure 5:**
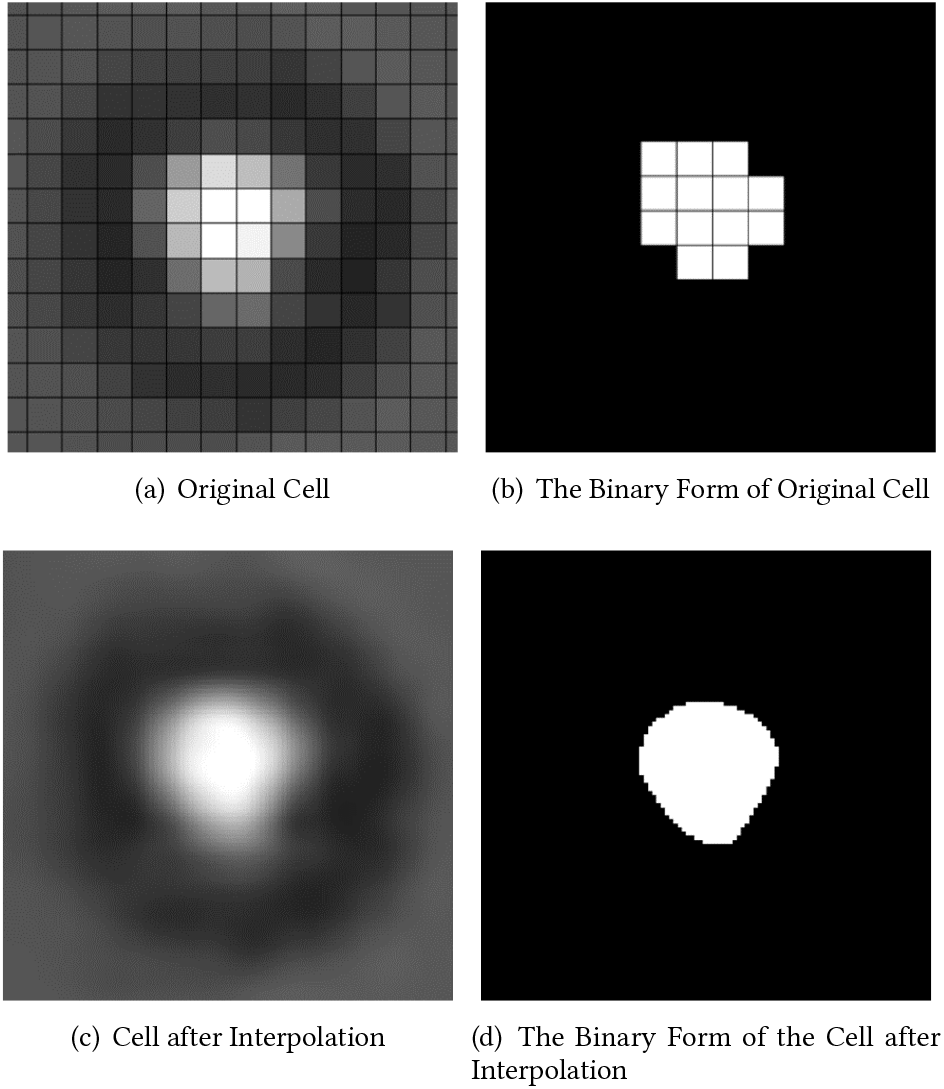
Images Scaling.

Figure 5 (c) is a rescaled version of Figure 5 (a) that used bicubic interpolation to enlarge it eight times. Figure 5 (d), the binary form of 5 (c), is generated using the same threshold used to get Figure 5 (b) from 5 (a). This rescaling smooths out the cell and portrays a more accurate shape demonstrated by comparing 5 (d) to the rest of the figures. Hence, for all further analyses, we will use the eight times enlarged images.

### 2.2 Cell Detection

In the original image, cells are the white spots surrounded by a black edge (cytomembrane), as shown in Figure 6 (a). Each image goes through 3 steps before we can detect a cell. First, the binary image is generated from the gray scale image using a pixel value threshold. In this paper, we set the threshold to 85 and convert all gray scale images into binary. The black pixels in Figure 6 (a) with a value below the preset threshold become converted into white pixels in Figure 6 (b). All other pixels above the threshold values convert to black pixels. This conversion is done by following the equation below:

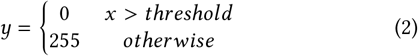

where *x* is the pixel value in the gray image, as is shown in Figure 6 (a), and *y* is the new pixel value in the binary image, as is shown in 6 (b).

**Figure 6:**
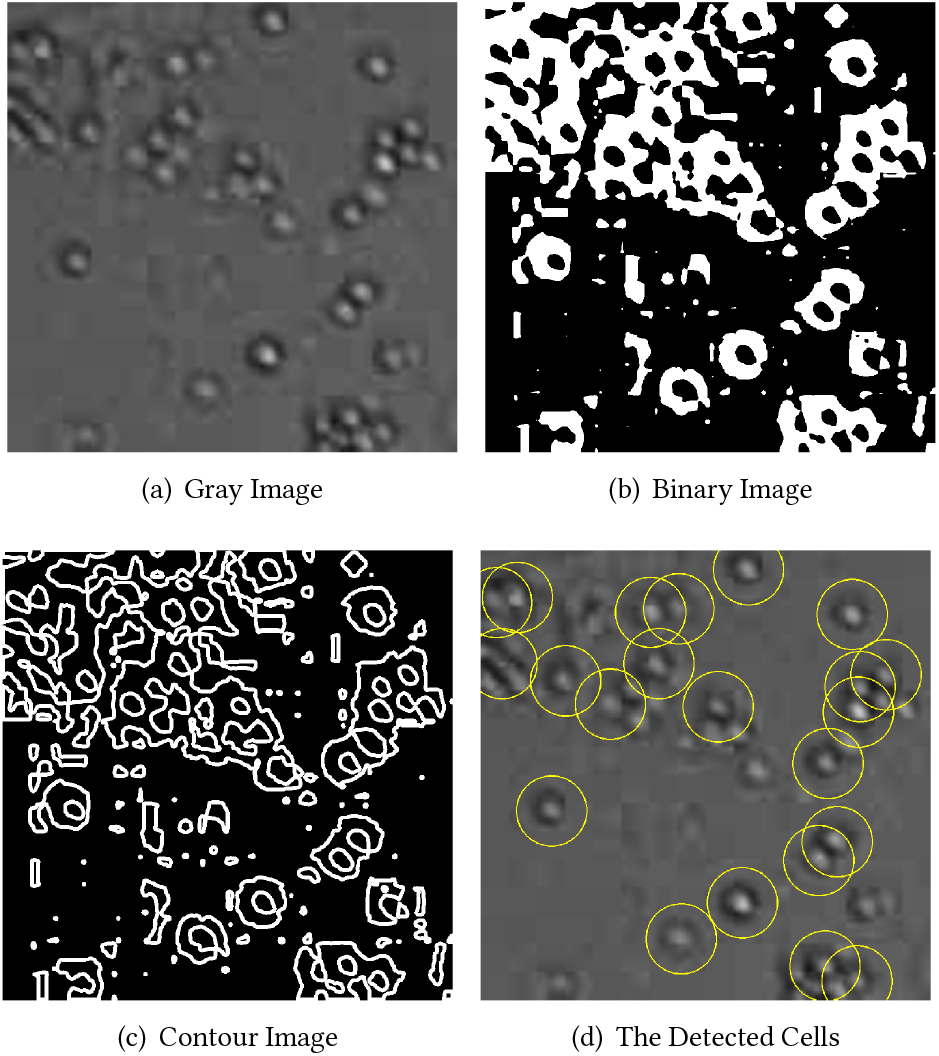
Cell Detection Steps.

Second, using the binary image, Figure 6 (b), we find the contours of the white pixels shown in Figure 6 (c) by applying the Border Following Contours Retrieving Algorithm [22]. Finally, the last step is to determine which contours represent a cell. Figure 6 (c) reveals that a cell typically has two contours, a larger and smaller one, which together forms an annulus. The annulus corresponds to the cytomembrane of the cell. In order to achieve a more accurate detection result, we use the following criterion to determine if an object is a cell:

1. A bigger contour must include a smaller one.
2. The radius of the smaller contour should be in a reasonable range. In this paper, the contour’s radius must be greater than 7 pixels and less than 90 pixels. We get the radius of a contour by using the smallest circle that can enclose that contour. The radius and center of this circle denote the radius and center of the contour.
3. We check if the pixels within the smaller contour is bright enough. In this paper the pixel value at the center should be larger than 100 to meet this condition.

If all three conditions are met, we conclude that the object is a cell. Figure 6 (d) displays all cells detected by our model from Figure 6 (a). We track the cells in the following steps using the coordinates of the center of the small contour to represent the coordinates of the cell.

### 2.3 Cell Tracking

Cell tracking across different frames (time points) is the next step in our framework. In this step, we map the detected cells in the current image to the detected cells in a previous image, shown in Figure 7. We create one track for each cell to record its position and shape across the time points.

**Figure 7:**
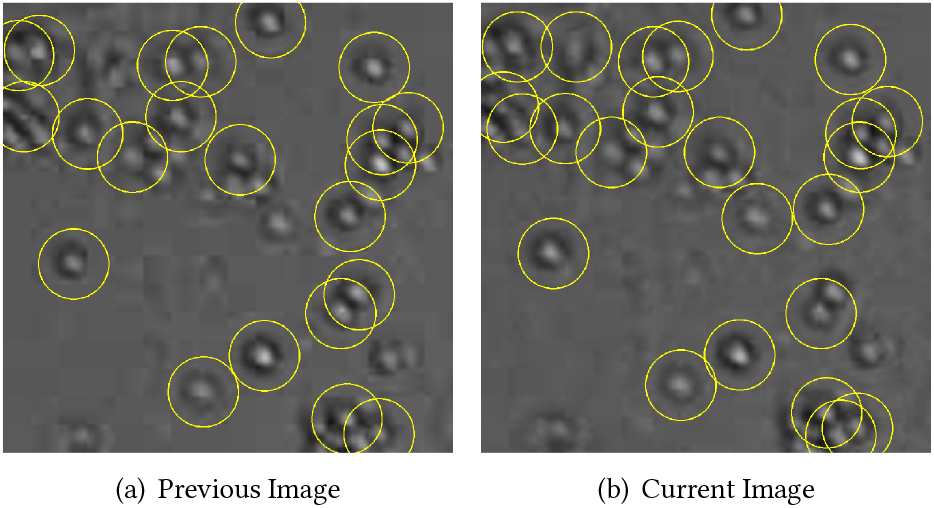
The Result of Cell Detection.

We use the Hungarian algorithm [8, 12] to find one to one mapping between cells of two images. Let *m* and *n* be the number of cells in our current and previous images, respectively. For each cell at an earlier time point, we superimpose it on the current image and calculate the distances to all *m* cells. Since we get the coordinates of all cells in the Cell Detection step, we can calculate the Euclidean distance between any pair of cells.

Figure 8 shows the current image. The detected cells are in the yellow circles, whereas the red circle denotes a cell’s position in the previous image. For every cell in the current image, the yellow circles, we calculate the distance between it and each cell in the previous image, the red circle. For *n* cells in the previous image and *m* cells in the current image, we end up with a *n × m* distance matrix.

**Figure 8:**
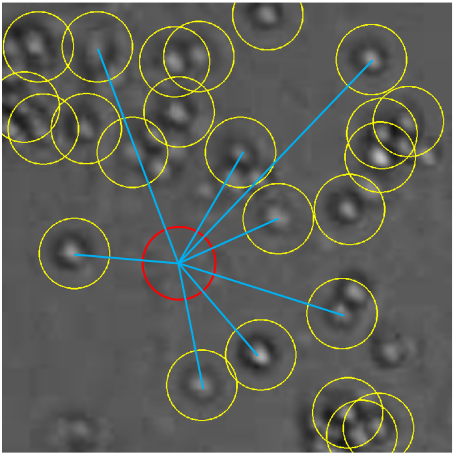
Cell Distances.

The Hungarian algorithm works based on the minimization of the distance matrix. The algorithm finds a value for each row such that the summation of all the values is minimum. If for the *i*^*th*^ row, the algorithm chooses a value in *j*^*th*^ column, it means that the *i*^*th*^ cell in the prior time point is the same as the *j*^*th*^ cell in the present time point. Applying the Hungarian algorithm on our matrix, we get the relation between all cells in the previous image and all cells in the current image. When *n* is not equal to *m*, some cells are unable to be matched. For example, *n* being bigger than *m* means there are more cells in the previous image; some of them have died or got devoured by macrophages, therefore, can not be tracked. If *n* is smaller than *m*, then there are more cells in the current image, so we need to create more tracks to record the movement of the extra cells accordingly.

As is shown in Figure 9, we mark the cells with a number in all-time points. If one cell matches between previous and current images, we use the same unique number of the prior time point to mark that cell in the present time point. This permits us to track the cell in the continuous images, allowing us to see how the cell changes through the time points.

**Figure 9:**
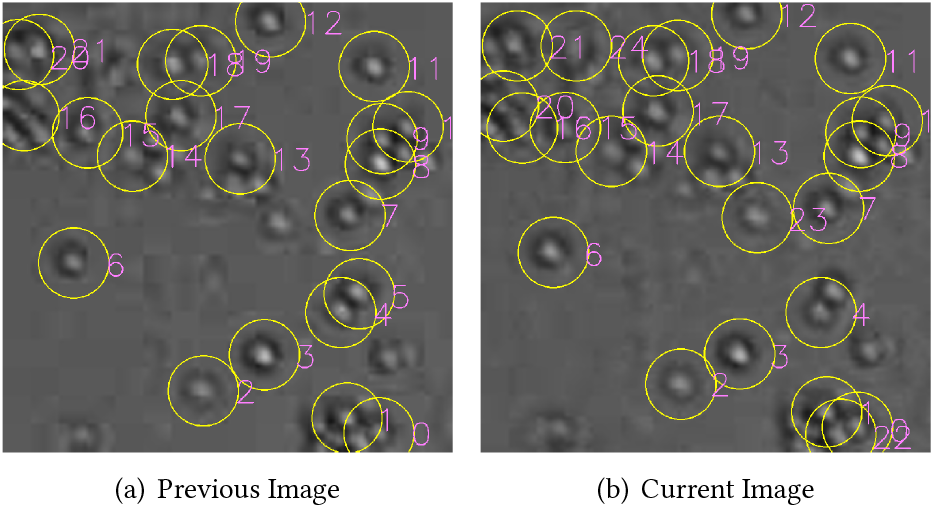
The Result of Cell Tracking.

### 2.4 Cell Classification

After we can successfully detect cells and track their position at each time point, we need to classify them between living and dead. To distinguish the live cells from the dead ones, we use the shape of the cytomembrane. It uses the fact that the shape of a live cell keeps changing over time (as shown in Figure 10), whereas the shape of a dead cell remains unchanged.

**Figure 10:**
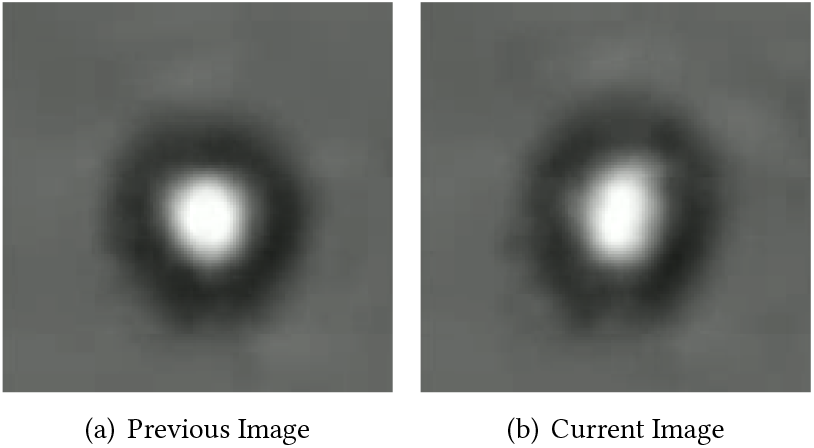
Cell Changes Shape.

To describe the shape of a cell, we use the minimum bounding box of the inner contour of that cell, shown in Figure 11. The minimum bounding box is the smallest rectangle that can encircle the inner contour of the cell, which is determined by the rotating calipers algorithm [18]. To track the cell changes over time, we propose a new metric, Ratio of Short side to Long side (RSL), which is defined by the following equation:

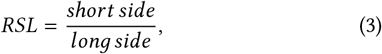

where RSL is the ratio of the shorter side of the rectangle to the longer side, as shown in Figure 11. A larger change in RSL value indicates a more substantial change in the shape of the cell, which directly relates to a cell being alive.

**Figure 11:**
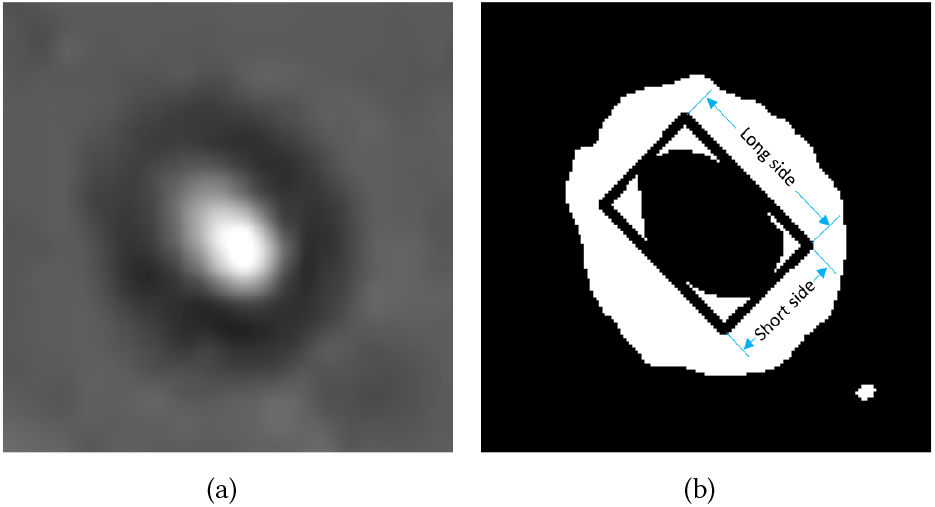
The Shape of One Cell.

To determine if a cell is alive at a given time point, we measure how much the cell shape changes across four previous and four following reference images. For example, if our given time point is 20 and we want to decide if a cell is alive at that moment, then we will consider time points 16-19 and 21-24 as reference. Figure 12 shows a sample of nine time points (eight references and one given). If the shape changes significantly between any two of the nine images then we can safely conclude that the cell is live at our assigned time point.

**Figure 12:**
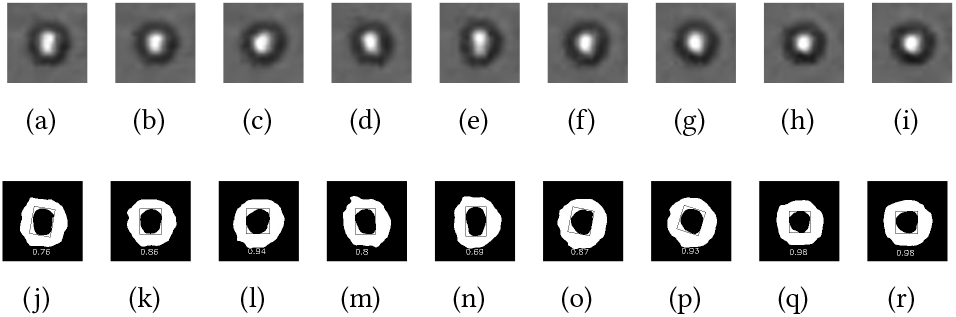
The change of RSL.

In Figure 12, (a-i) are the original cells and (j-r) illustrate their binary form. The nine RSLs are 0.76, 0.86, 0.94, 0.8, 0.69, 0.87, 0.93, 0.98, 0.98 respectively. The maximum among the nine RSLs is 0.95 and the minimum is 0.63. The difference between the maximum and minimum represents the maximum change of shape of this cell between any of the nine time points. We set a threshold of 0.3 for change in RSL to reduce computational uncertainties. For this example, the maximum change in RSL is 0.32 which is above the threshold. Therefore, we can conclude that this cell is live at a given time point (represented by Figure 12 (e)). Any value smaller than 0.3 for other cases will indicate that the cell is dead.

### 2.5 Detection of Phagocytosis

As previously mentioned, a macrophage is a type of phagocyte that detects, engulfs, and destroys pathogens like a tumor cell. This event would cause the number of cells to decrease locally. We can identify phagocytosis events by clustering the cells based on proximity or density and tracking the changes in the number of cells in clusters. If cells get devoured in a particular cluster, we will see a significant drop in cell population of that cluster across different time points. In this paper, we use the DBSCAN algorithm [7] to cluster cells, as is shown in Figure 13, where the number beside each cluster represents the cluster index.

**Figure 13:**
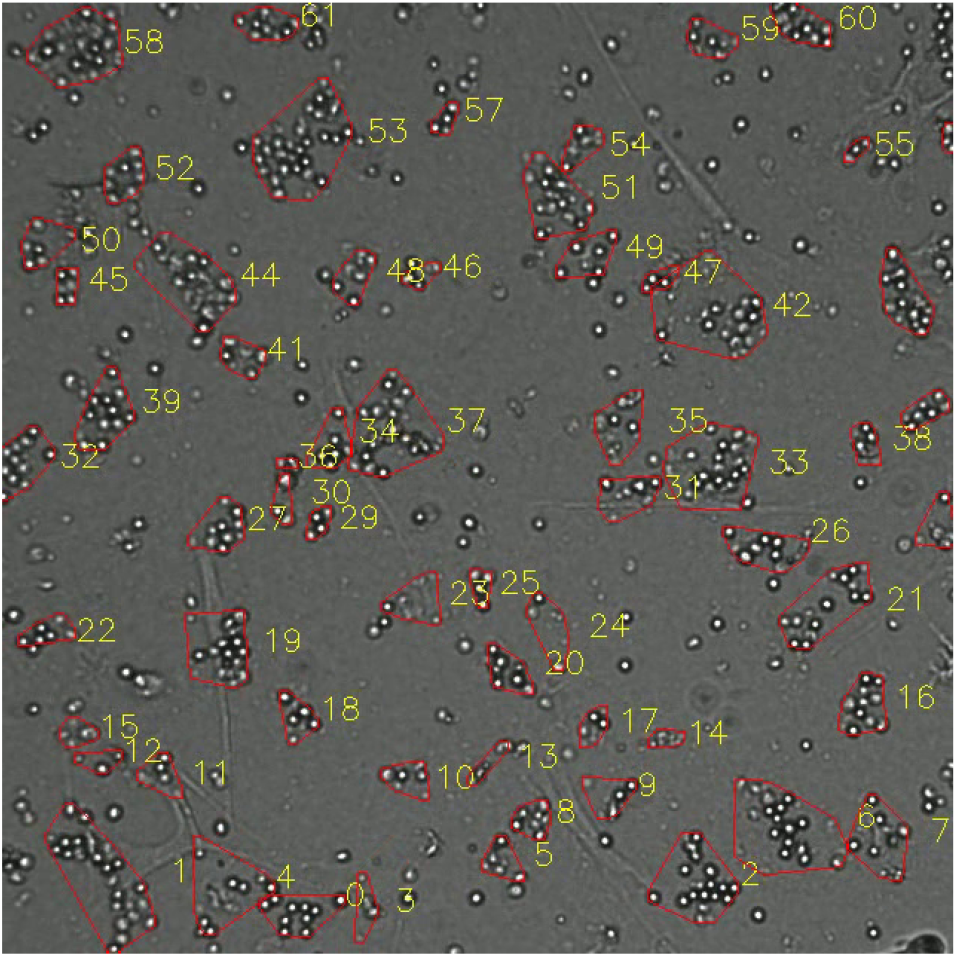
Cell Clusters.

As the cells are continuously moving, cluster structures will change even without any phagocytosis. Some cells may move further from a cluster resulting in it being excluded from that cluster. At the same time, it will be included to a new cluster based on density, as can be seen in Figure 14. In Figure 14 (a), the cell pointed by the yellow arrow moves closer to the cluster so that the cluster includes it. On the contrary, Figure 14 (b) illustrates a cell marked by a yellow arrow that is moving far from the cluster; consequently, the cluster will then exclude the cell. Phagocytosis will cause much faster decreases in the cluster population than random cell movement alone. We need to set a threshold for change in the cluster population to differentiate between these two cases.

**Figure 14:**
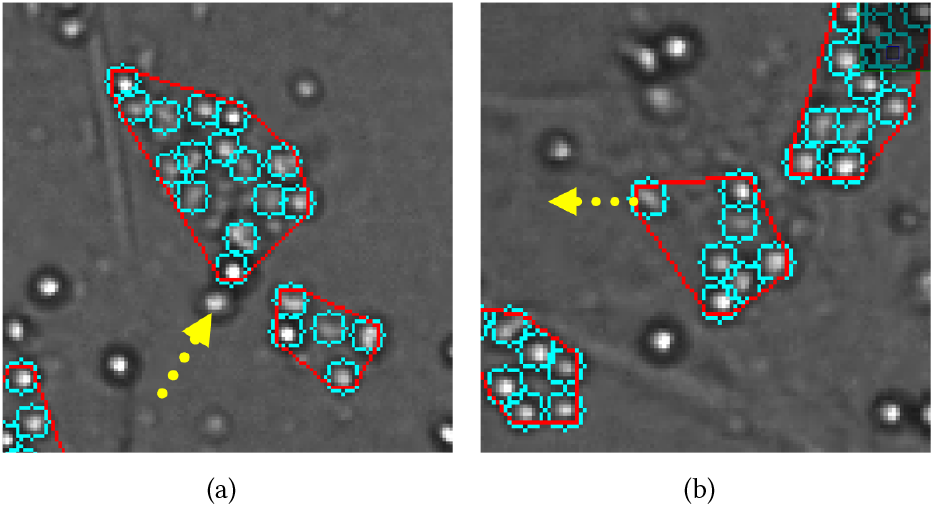
The Change of the Clusters.

When phagocytosis happens in a cluster, many cells disappear as time goes on. Figure 15 illustrate this phenomenon. The cells in the red circle make up Cluster 53. The cell in the blue box is the macrophage. There are 29, 18, 9 cells within Custer 53 in Figure 15 (a) (b) (c), respectively. The macrophage in this cluster is very active and it engulfs cancer cells when they are in contact with each other.

**Figure 15:**
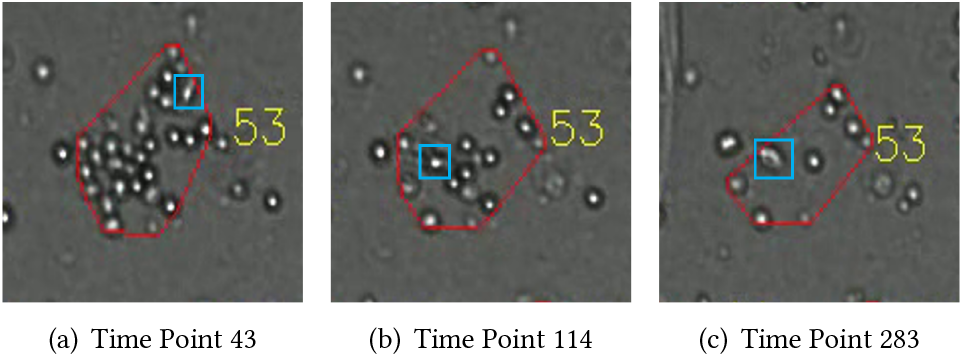
The Number of Cells Decreases in Cluster 53.

We count the number of cells in every time point and check if the number within a cluster decreases. In Figure 16, we use linear regression to fit a straight line to show the rate of reduction of the cell population in Cluster 53. If the slope of the fit line is steeper than −0.04, we will conclude that the cluster contains phagocytosis. The slope of this fit line for Cluster 53 is −0.086, which surpasses the preset threshold. Accordingly, we can conclude that Cluster 53 contains phagocytosis.

**Figure 16:**
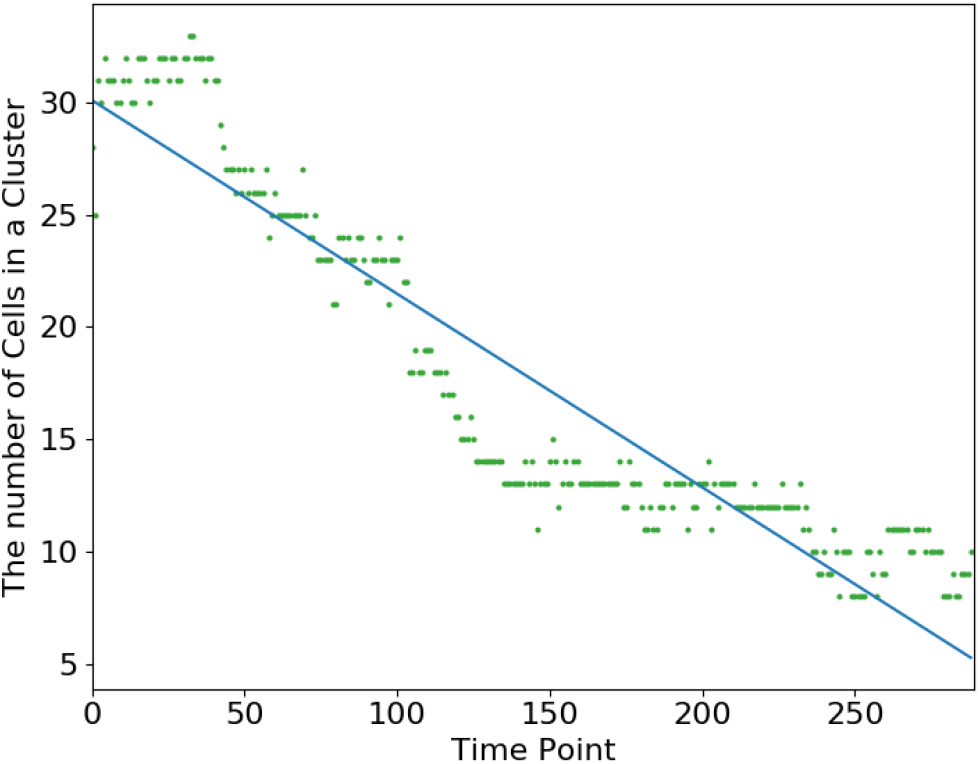
The Number of Cells Changes in Cluster 53.

## 3 EXPERIMENTS

In this section, we performed two experiments to evaluate the framework. First, we tested the performance of our classification model using a simulated video that contains both live and dead cells with 193 time points. Furthermore, we added different levels of noise to that simulated video to test the robustness of our framework. Second, we test our framework for its detection of phagocytosis. In this experiment, we used two time-lapse microscopy imaging data (‘videos’) as the input to detect the events.

### 3.1 Classification of Live and Dead Cells

The cells in original images are not labeled, which means we do not have any ground truth to quantify the accuracy of our classification. Our collaborators manually labeled nine cells from the original imaging data and classified them into three categories: always live, live to dead, and always dead. We simulated a series of images (193 time points) using the nine labeled cells. Each of the nine cells were duplicated 20 times in each image. Among the 20 duplicates, each quarter were rotated 0, 90, 180, 270 degrees, respectively, resulting in four different orientations for one cell. In the original imaging data, if any of the nine labeled cells disappears at any time point, all of its duplicates will also disappear at the same time point in the simulated data. In the first simulated time point, the cells were arbitrarily positioned as long as they did not overlap. In the following time points they were given random movements.

We applied our algorithm to classify the cells into living and dead and yielded 100% accuracy for the simulated cells at all time points. As for the robustness of our algorithm, we added three different levels of Gaussian noises *N* (*μ*, *σ*^2^) into the background of the simulated images, where *μ* = 0 and *σ* = 0.5, 0.6, and 0.7 correspondingly. The larger the *σ* (standard deviation), the more random noise we added to the image background. The detection result with the noise can be seen in Figure 17. The horizontal axis represents the time points and the vertical axis represents classification accuracy. Accuracy is the ratio of the number of correctly classified cells to the total number of cells in the experiment. From Figure 17, we can see that the accuracy of our algorithm is higher than 0.8, even with the largest background noise (*σ* = 0.7) we added. It indicates that the cell detection, tracking, and classification steps are robust to the background noises in the real time-lapse microscopy imaging data.

**Figure 17:**
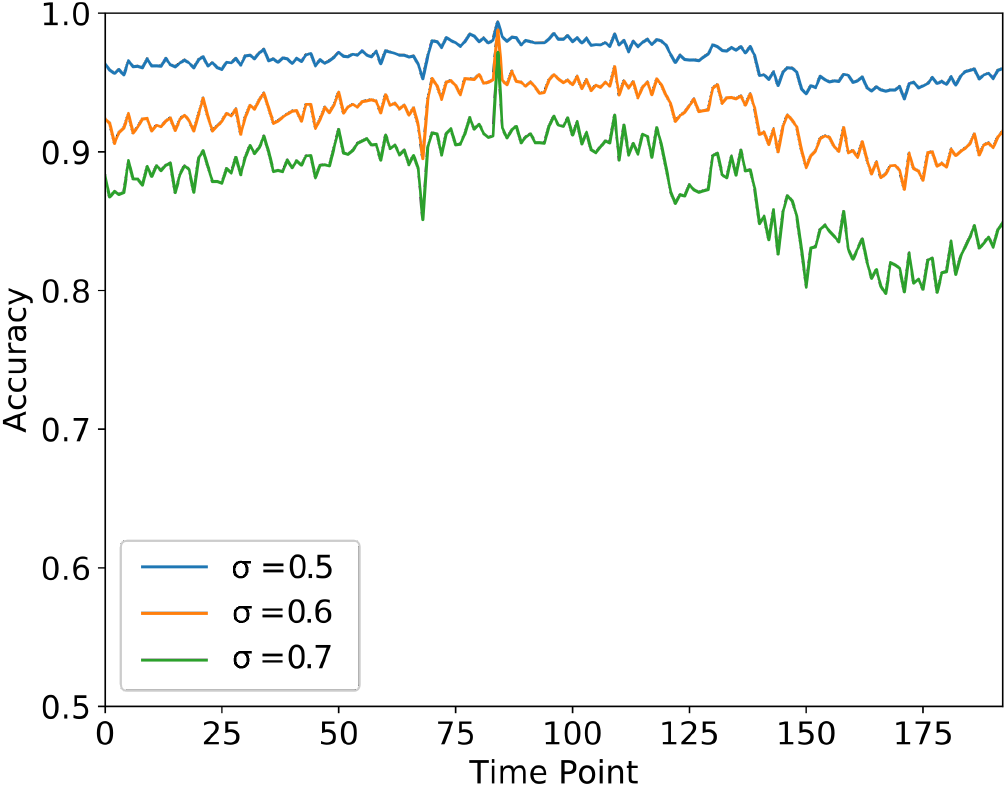
Classification Accuracy with Different Background Noise.

### 3.2 Detection of phagocytosis

We processed two real time-lapse microscopy imaging data (videos) to check whether they have phagocytosis in any cluster of cells. The result from the first video (one time point) is shown in Figure 18. In Figure 18 (a), the clusters that are determined to have phagocytosis are circled by red, whereas the other clusters are encircled by yellow. The number beside each cluster represents its index in that image. Figure 18 (b) shows the change in the number of cells in three selected clusters. The number of cells in Clusters 117 and 137 decrease as time goes by and the slopes are higher than the threshold, which indicates that clusters have phagocytosis. Cell populations in cluster 121 fluctuate, which is attributed to the random movement of cells.

**Figure 18:**
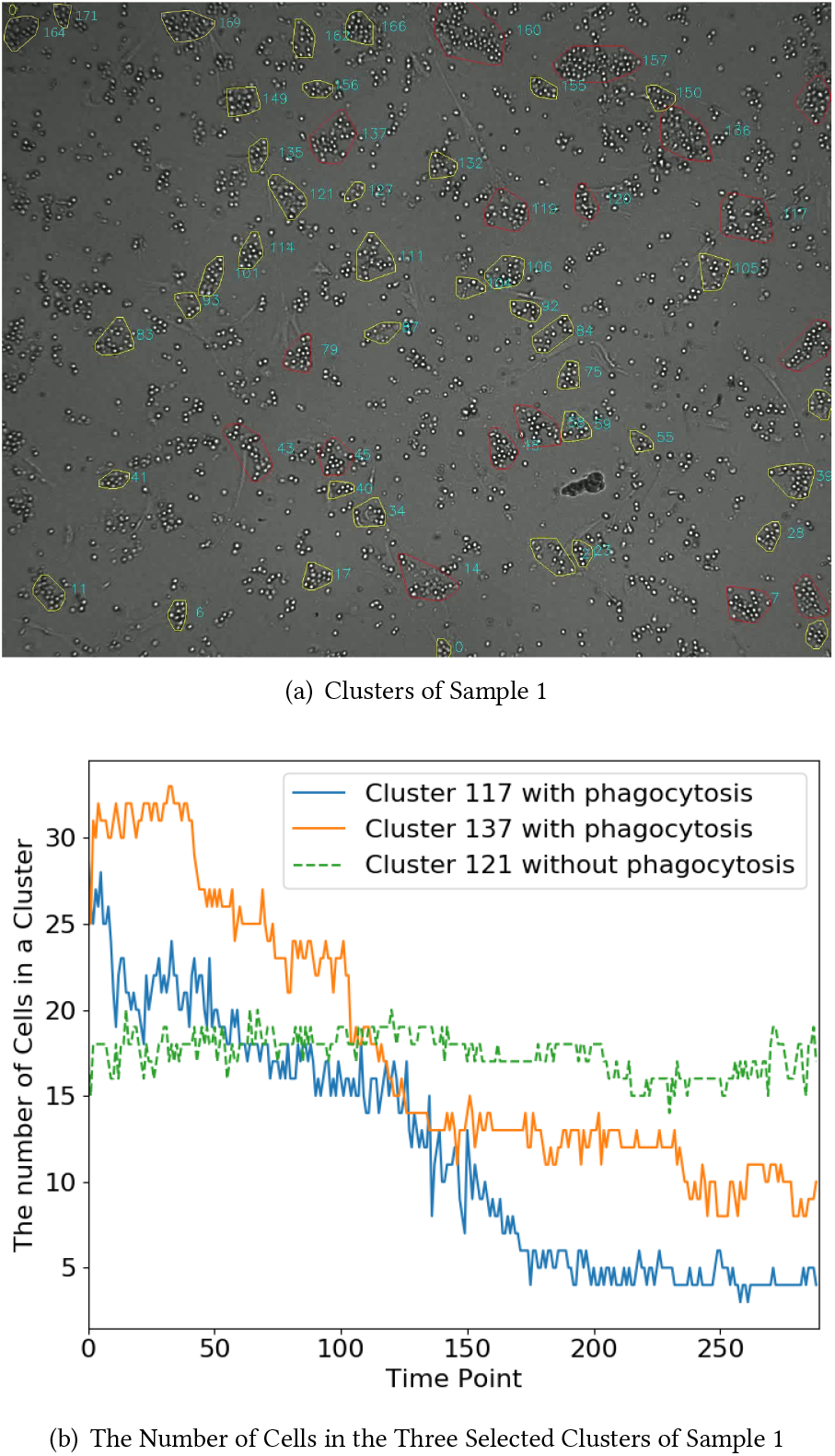
Clusters in Sample 1.

The result from the second video (one time point) can be seen in Figure 19. Three clusters were selected in the video to show the change of cell numbers within the clusters. We can see that Clusters 23 and 83 in Figure 19 (b) have a stable decrease in number of cells. As explained in the Methods Section, we use linear regression to model the decreasing trend. The slopes of the fitted regression lines for the number of cells in the two clusters are above the threshold, indicating the presence of phagocytosis. The fitted line of Cluster 41 in 19 (b) is stable in a range without any apparent decreasing trend. Therefore, we can classify it into the category of clusters that has no phagocytosis.

**Figure 19:**
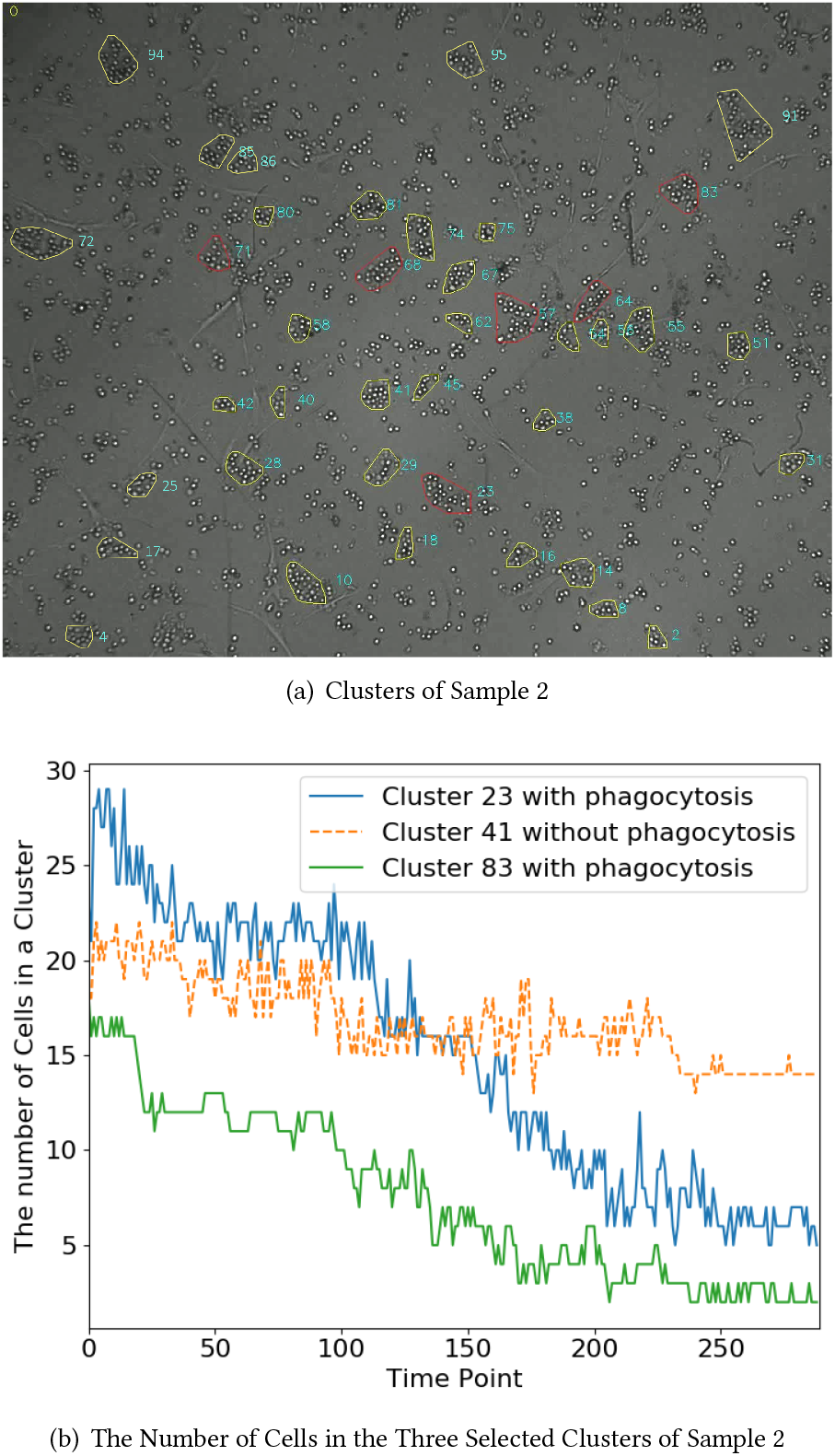
Clusters in Sample 2.

In each video, several clusters are determined to have phagocytosis that is illustrated by the clusters circled by red in Figures 18 and 19. The results can show that the cells within these clusters are devoured by macrophage gradually, which demonstrates that the proposed method is very effective. The pathologists confirm most of these detected phagocytosis.

### 3.3 Running Time

To measure the scalability of the proposed framework, we tested it on a 289-image time-lapse microscopy data. The framework took 180 CPU seconds to detect, track, and classify all the cells in one experiment. The CPU time was measured on AMD Ryzen Threadripper 2950X CPU with 2140MHz.

## 4 DISCUSSION AND CONCLUSION

Time-lapse microscopy imaging data estimates the response of tumor samples to anti-cancer drugs at single-cell resolution, which enables cell tracking and the observation of a variety of cellular activities to study anti-cancer therapeutics. In this study, we propose an efficient framework for analyzing the large-scale time-lapse microscopy imaging data to track the behaviors of thousands of cancer cells simultaneously. The overall framework can be divided into five main steps. For each step, we describe the process, math formulas, and its importance. In the first and second steps, an advanced image interpolation technique is applied to improve the image resolution. Then, the cell contours are automatically estimated for cell detection. In the third step, the Hungarian algorithm is applied to track cells among the images at different time points in the same experiment. It calculates the pairwise distances of cells between two consecutive time points. In some studies [1, 13], Kalman filtering [9] was used to track multiple moving cells and estimate the path of each cell. However, cell migration is usually considered as Brownian motion [24] and the trajectories of the cells are difficult to predict. Therefore, the Hungarian algorithm is not integrated with the Kalman filter in this study. In the fourth step, we classify the live and dead cells based on whether the cells undergo deformations at different time points. If yes, we classify it as a live cell. In the last step, we apply a density-based clustering algorithm (i.e., DBSCAN) to detect phagocytosis events.

The results in the simulation experiment demonstrate that the proposed framework is robust to image background noise and it can accurately track and classify the live and dead cells in each image. The experiments on detecting phagocytosis events illustrate that the pipeline has great potential to identify macrophages and track their behaviors. Overall, the work in this paper introduces a comprehensive pipeline for time-lapse microscopy imaging data analysis and provides an accurate estimation of the drug response for each cancer patient.

## ACKNOWLEDGEMENT

This research work is supported by H. Lee Moffitt Cancer Center & Research Institute Foundation. Tara Rutkowski is supported by National Science Foundation Research Experiences for Undergraduates (REU) Program under project IIS 1755761.

## REFERENCES

[1] Badarish Colathur Arvind, Sujith Kumar Nagaraj, Chandra Sekhar Seelamantula, and Sai Siva Gorthi. 2016. Active-disc-based Kalman filter technique for tracking of blood cells in microfluidic channels. In 2016 IEEE International Conference on Image Processing (ICIP). IEEE, 3394–3398.

[2] Mark-Anthony Bray and Anne E Carpenter. 2015. CellProfiler Tracer: exploring and validating high-throughput, time-lapse microscopy image data. BMC bioinformatics 16, 1 (2015), 369.

[3] Felix Buggenthin, Carsten Marr, Michael Schwarzfischer, Philipp S Hoppe, Oliver Hilsenbeck, Timm Schroeder, and Fabian J Theis. 2013. An automatic method for robust and fast cell detection in bright field images from high-throughput microscopy. BMC bioinformatics 14, 1 (2013), 297.

[4] Joe Chalfoun, Michael Majurski, Alden Dima, Michael Halter, Kiran Bhadriraju, and Mary Brady. 2016. Lineage mapper: A versatile cell and particle tracker. Scientific reports 6 (2016), 36984.

[5] Oleh Dzyubachyk, Wiro Niessen, and Erik Meijering. 2008. Advanced levelset based multiple-cell segmentation and tracking in time-lapse fluorescence microscopy images. In 2008 5th IEEE International Symposium on Biomedical Imaging: From Nano to Macro. IEEE, 185–188.

[6] Oleh Dzyubachyk, Wiggert A Van Cappellen, Jeroen Essers, Wiro J Niessen, and Erik Meijering. 2010. Advanced level-set-based cell tracking in time-lapse fluorescence microscopy. IEEE transactions on medical imaging 29, 3 (2010), 852–867.

[7] Martin Ester, Hans-Peter Kriegel, Jörg Sander, Xiaowei Xu, et al. 1996. A density-based algorithm for discovering clusters in large spatial databases with noise.. In Kdd, Vol. 96. 226–231.

[8] Khuloud Jaqaman, Dinah Loerke, Marcel Mettlen, Hirotaka Kuwata, Sergio Grinstein, Sandra L Schmid, and Gaudenz Danuser. 2008. Robust single-particle tracking in live-cell time-lapse sequences. Nature methods 5, 8 (2008), 695.

[9] Rudolph Emil Kalman. 1960. A new approach to linear filtering and prediction problems. (1960).

[10] Takeo Kanade, Zhaozheng Yin, Ryoma Bise, Seungil Huh, Sungeun Eom, Michael F Sandbothe, and Mei Chen. 2011. Cell image analysis: Algorithms, system and applications. In 2011 IEEE Workshop on Applications of Computer Vision (WACV). IEEE, 374–381.

[11] Harpreet Kaur and Neelofar Sohi. 2017. A study for applications of histogram in image enhancement. Int. J. Eng. Sci 6, 6 (2017), 59–63.

[12] Harold W Kuhn. 1955. The Hungarian method for the assignment problem. Naval research logistics quarterly 2, 1–2 (1955), 83–97.

[13] Min Liu, Yue He, Yangliu Wei, and Peng Xiang. 2017. Plant cell tracking using Kalman filter based local graph matching. Image and Vision Computing 60 (2017), 154–161.

[14] Firas Mualla, Simon Schöll, Björn Sommerfeldt, Andreas Maier, and Joachim Hornegger. 2013. Automatic cell detection in bright-field microscope images using SIFT, random forests, and hierarchical clustering. IEEE transactions on medical imaging 32, 12 (2013), 2274–2286.

[15] Dale Muzzey and Alexander van Oudenaarden. 2009. Quantitative time-lapse fluorescence microscopy in single cells. Annual Review of Cell and Developmental 25 (2009), 301–327.

[16] Adrienne HK Roeder, Alexandre Cunha, Michael C Burl, and Elliot M Meyerowitz. 2012. A computational image analysis glossary for biologists. Development 139, 17 (2012), 3071–3080.

[17] S Sarkar, N Cohen, P Sabhachandani, and T Konry. 2015. Phenotypic drug profiling in droplet microfluidics for better targeting of drug-resistant tumors. Lab on a Chip 15, 23 (2015), 4441–4450.

[18] MI Shamos. 1978. Computational geometry[Ph. D. Thesis]. (1978), 76–-82.

[19] Ariosto Silva, Timothy Jacobson, Mark Meads, Allison Distler, and Kenneth Shain. 2015. An organotypic high throughput system for characterization of drug sensitivity of primary multiple myeloma cells. JoVE (Journal of Visualized Experiments) 101 (2015), e53070.

[20] Ariosto Silva, Maria C Silva, Praneeth Sudalagunta, Allison Distler, Timothy Jacobson, Aunshka Collins, Tuan Nguyen, Jinming Song, Dung-Tsa Chen, Lu Chen, et al. 2017. An ex vivo platform for the prediction of clinical response in multiple myeloma. Cancer research 77, 12 (2017), 3336–3351.

[21] Praneeth Sudalagunta, Maria C Silva, Rafael R Canevarolo, Raghunandan Reddy Alugubelli, Gabriel DeAvila, Alexandre Tungesvik, Lia Perez, Robert Gatenby, Robert Gillies, Rachid Baz, et al. 2020. A pharmacodynamic model of clinical synergy in multiple myeloma. EBioMedicine 54 (2020), 102716.

[22] Satoshi Suzuki et al. 1985. Topological structural analysis of digitized binary images by border following. Computer vision, graphics, and image processing 30, 1 (1985), 32–46.

[23] Philippe Thévenaz, Thierry Blu, and Michael Unser. [n.d.]. Image interpolation and resampling. ([n. d.]).

[24] Roumen Tsekov and Marga C Lensen. 2013. Brownian motion and the temperament of living cells. Chinese Physics Letters 30, 7 (2013), 070501.

[25] Leonardo F Urbano, Puneet Masson, Matthew VerMilyea, and Moshe Kam. 2016. Automatic tracking and motility analysis of human sperm in time-lapse images. IEEE transactions on medical imaging 36, 3 (2016), 792–801.

[26] Viktoria von Manstein, Chul Min Yang, Diane Richter, Natalia Delis, Vida Vafaizadeh, and Bernd Groner. 2013. Resistance of cancer cells to targeted therapies through the activation of compensating signaling loops. Current signal transduction therapy 8, 3 (2013), 193–202.

[27] N Ezgi Wood and Andreas Doncic. 2019. A fully-automated, robust, and versatile algorithm for long-term budding yeast segmentation and tracking. PloS one 14, 3 (2019).

